# *SYNGAP1* haploinsufficiency disrupts early neurodevelopment and accelerates intrinsic neuronal maturation in human patient-derived models

**DOI:** 10.64898/2026.07.15.738667

**Authors:** Montanna Waters, Lucas Teasdale, Sean Byars, Cristiana Mattei, Nessia Eve Roseno, Dmitry Ovchinnikov, Ingrid E Scheffer, Heath R Pardoe, Steven Petrou, Snezana Maljevic

**Author notes:** **Corresponding Author:** Associate Professor Snezana Maljevic, The Florey Institute of Neuroscience and Mental Health 3052 Parkville, Victoria, Australia, **E-mail:**. These authors contributed equally to this work and share first authorship (Montanna Waters, Lucas Teasdale).

## Abstract

*SYNGAP1* developmental and epileptic encephalopathy (DEE) is a severe neurodevelopmental disorder characterised by intellectual disability, developmental delay, and refractory epilepsy caused by heterozygous variants in *SYNGAP1*, which encodes Synaptic Ras GTPase-activating protein 1. While *SYNGAP1* is best known for its role at the postsynaptic density, increasing evidence indicates that haploinsufficiency also disrupts early neurodevelopment. Here, we used patient-derived induced pluripotent stem cell (iPSC) models to investigate early neurodevelopmental and neuronal phenotypes associated with *SYNGAP1* haploinsufficiency. iPSCs derived from a female patient carrying the frameshift variant p.Leu150Valfs*6 were differentiated into two complementary models: micropatterned neural rosettes representing early neuroepithelial organisation and NGN2-induced excitatory neurons representing postmitotic functional development. Patient-derived neural rosettes displayed enlarged, dysmorphic lumens, indicating disrupted neuroepithelial organisation at the earliest stages of brain development. Transcriptomic profiling revealed widespread dysregulation of genes involved in neurodevelopment, cell adhesion and ion channel regulation, including coordinated downregulation of protocadherin family members. Whole-cell patch-clamp electrophysiology demonstrated reduced input resistance, larger action potential amplitudes, and increased inward and outward current densities, consistent with accelerated intrinsic neuronal maturation rather than generalized hyperexcitability. Together, these complementary findings demonstrate that *SYNGAP1* haploinsufficiency disrupts early human brain development and accelerates intrinsic neuronal maturation, with pathogenic mechanisms emerging before synaptogenesis and extending beyond *SYNGAP1*’s established synaptic role.

## Introduction

*SYNGAP1* developmental and epileptic encephalopathy (DEE) is a severe neurodevelopmental disorder characterized by intellectual disability, developmental delay, seizures, and autism spectrum disorder ^1,2^. *SYNGAP1* variants are a significant contributor to intellectual disability, accounting for approximately 1-2% of cases worldwide ^3,4^. *SYNGAP1* plays a crucial role in the postsynaptic density, where it associates with the NMDA receptor complex and regulates the ERK/MAPK pathway signalling following calcium influx ^5^. Although much of the research on *SYNGAP1* DEE has focused on synaptic dysfunction and seizure mechanisms, increasing evidence suggests that *SYNGAP1* also plays an important role in early neurodevelopment.

Homozygous knockout of *SYNGAP1* is embryonically lethal in mice ^5^, demonstrating an essential role during development. Haploinsufficiency of *SYNGAP1* accelerates maturation of excitatory synapses in the mouse neocortex, restricting the window of synaptic plasticity and impairing cortical circuit assembly ^6^. In addition, *SYNGAP1* haploinsufficiency disrupts the structural integrity of pyramidal cells, impacting dendritic growth and spine plasticity ^7^. Together, these findings suggest that *SYNGAP1* variants may influence neurodevelopmental processes before the establishment of mature synaptic networks. Consistent with this, recent work by Birtele et al. ^8^ demonstrated that *SYNGAP1* is expressed in radial glial cells and regulates cytoskeletal dynamics, cortical lamination, and accelerated maturation of cortical projection neurons.

Human induced pluripotent stem cell (iPSC)-derived neural models provide a powerful platform for investigating disease-associated developmental mechanisms while preserving the patient’s genetic background. Among these, NGN2-induced excitatory neurons provide a rapid and highly reproducible system for analysing transcriptional and electrophysiological phenotypes. However, these approaches are limited in their ability to recapitulate the earliest stages of brain development, including neurulation, cell differentiation and tissue patterning. To address these limitations, recent advances have focused on developing reproducible models that capture early human brain development.

Knight and colleagues ^9^ developed a geometrically restricted micropatterned neural rosette system that recapitulates key features of human neuroepithelial organisation and has already revealed morphological abnormalities in *SYNGAP1* patient-derived iPSCs ^8^. Neural rosettes comprise polarised neural progenitor cells surrounding a central lumen and reproduce key structural and cellular features of the developing neural tube during early human brain development ^10^. Their relatively simple cellular composition and sensitivity to developmental perturbation make them a valuable model for investigating early neurodevelopmental disorders^9^.

Here, we investigated the developmental consequences of *SYNGAP1* haploinsufficiency using patient-derived iPSCs carrying the pathogenic frameshift variant p.Leu150Valfs*6. We combined micropatterned neural rosettes to assess early neuroepithelial organisation with NGN2-induced excitatory neurons to examine transcriptional alterations and intrinsic electrophysiological maturation. Together, these complementary models enabled us to define developmental abnormalities that emerge before synaptogenesis and extend beyond *SYNGAP1*’s established synaptic role.

## Results

### Molecular characterisation of *SYNGAP1* iPSC lines

In this study we employed an iPSC line obtained from fibroblasts of a female patient harbouring the c.435_447dup mutation, a 13-bp duplication resulting in a leucine-to-valine substitution at position 150 and a frameshift causing premature termination (p.Leu150Valfs*6), also identified in the patient’s sister ^2^ (Fig. 1A). Successful reprogramming and retention of the variant were confirmed by PCR and Sanger sequencing using primers flanking the duplication site. Patient iPSCs displayed a double band on gel electrophoresis, consistent with heterozygous carriage of the duplication, while the healthy female control line (WC026i-5807-3) produced a single band at 241 bp (Fig. 1B; Supplementary Figure S1A). Patient and control iPSC lines also expressed the pluripotency markers NANOG and TRA-1-60, confirming maintenance of pluripotency (Supplementary Figure S1B).

**Figure 1.**
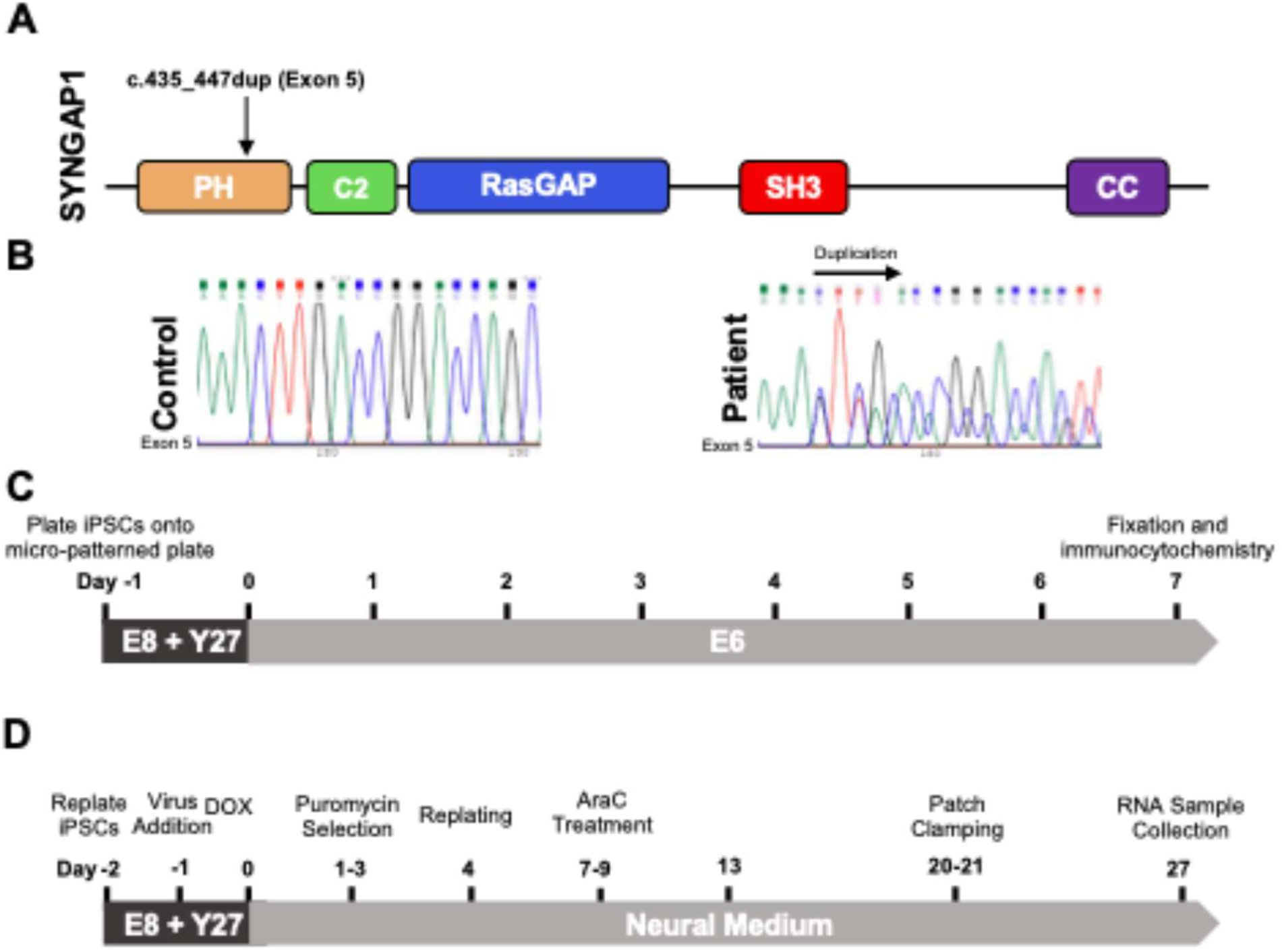
Characterisation of *SYNGAP1* c.435_447dup iPSC-derived neural cultures. (A) Schematic representation of the *SYNGAP1* protein showing major functional domains (PH, C2, RasGAP, SH3, CC) and the location of the patient variant (c.435_447dup, pLeu150Valfs*6). (B) Sanger sequencing traces from control and patient iPSCs. The patient carries a heterozygous duplication on exon 5, indicated by the peak shift at the duplicated region (arrow). (C) Experimental timeline for differentiation of iPSCs into neural rosettes. iPSCs were replated onto micropatterned plates on DIV −1. From DIV 0, cultures were maintained in E6 medium to initiate neural fate before fixation and immunocytochemistry at DIV 7. (D) Experimental timeline for differentiation of iPSCs into NGN2 neural cultures. iPSCs were replated on DIV −2, followed by lentiviral transduction and doxycycline induction on DIV −1 to 0. Puromycin selection occurred on DIV 1–3, and cells were replated on DIV 4. AraC treatment was performed on DIV 7–9 to limit proliferating cells, and cultures were maintained in neural medium from DIV 0 onward. Patch-clamp recordings were performed at DIV 20–21, and RNA was collected at DIV 27.

To examine whether *SYNGAP1* haploinsufficiency disrupts early neurodevelopment, we differentiated patient and control iPSCs into two complementary models: micropatterned neural rosettes using the Knight and Lundin ^9^ protocol to capture early neuroepithelial organisation (Fig. 1C), and NGN2-induced excitatory cortical neurons to assess cell-autonomous transcriptional and electrophysiological phenotypes (Fig. 1D, Supplementary Figure S1C).

### *SYNGAP1* iPSC-derived neural rosettes exhibit enlarged and dysmorphic lumens

Micropatterned neural rosettes were generated from patient and control iPSCs using the Knight and Lundin protocol ^9^, producing geometrically constrained neuroepithelial structures that recapitulate early neural tube organisation. Rosettes were immunostained for PAX6 (neural progenitor identity) and N-cadherin (apical cell adhesion), with DAPI counterstaining, and imaged by spinning disk confocal microscopy. Lumen morphology was quantified from z-projected images using semi-automated image analysis. Measurements were derived from 328 control and 718 patient rosettes pooled across three independent differentiations and two patient iPSC clones.

Immunostaining for PAX6 and N-cadherin revealed polarized radial neuroepithelial structures in both control and *SYNGAP1* DEE cultures, with apparent differences in lumen size and shape (Fig. 2A). Quantitative morphometric analysis confirmed these observations (Fig. 2B–G). Overall rosette area (40283 ± 140.8 vs 38063 ± 179.6 µm^2^; Fig. 2B, P > 0.05) and circularity (0.8789 ± 0.0011 vs 0.8749 ± 0.0007; Fig. 2C, P > 0.05) were comparable between genotypes, indicating that overall rosette morphology was preserved. In contrast, *SYNGAP1* DEE rosettes exhibited significantly fewer lumens per rosette (0.70 ± 0.021 vs 0.92 ± 0.040; Fig. 2D, P < 0.0001), enlarged lumen area (6793 ± 105.6 vs 5347 ± 211.6 µm^2^; Fig. 2E, P < 0.0001), reduced lumen circularity (0.7211 ± 0.005 vs 0.7678 ± 0.0053; Fig. 2F, P < 0.0001), and lumens positioned significantly closer to the rosette centre (14.6 ± 0.46 vs 23.62 ± 1.15 µm; Fig. 2G, P < 0.0001). Together, these findings indicate that the structural disruption is specific to the apical lumen rather than reflecting a general difference in rosette size or morphology.

**Figure 2.**
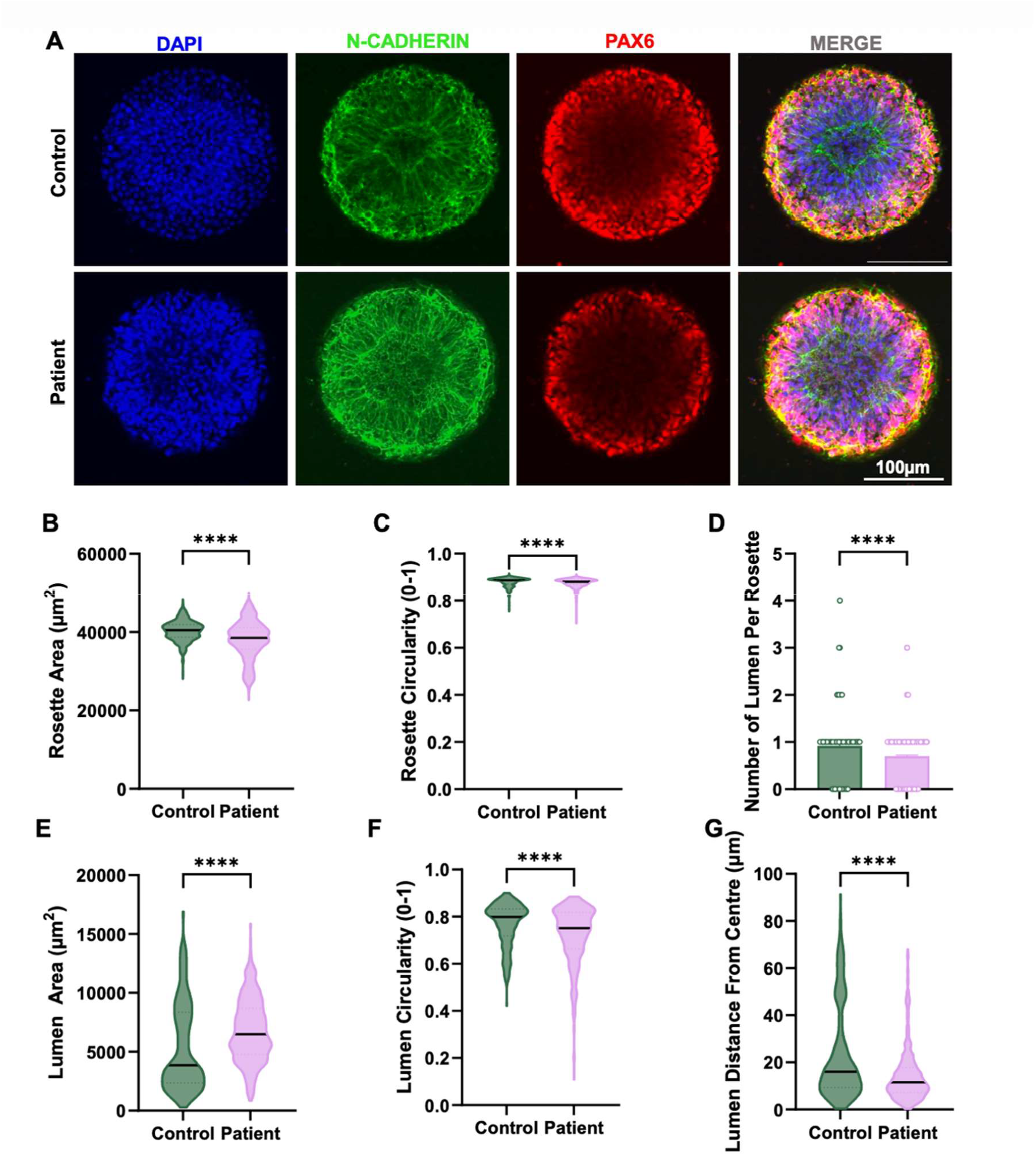
Neural rosettes derived from *SYNGAP1* DEE patient iPSCs exhibit enlarged and dysmorphic lumens. (A) Representative immunofluorescence images of single neural rosettes generated from control and *SYNGAP1* DEE patient iPSCs. Nuclei are labelled with DAPI (blue), the apical lumen with N-cadherin (green), and neural progenitors with PAX6 (red). Scale bar, 100 µm. (B–G) Quantitative analysis of neural rosette morphology. Measurements include (B) rosette area (control 40283 ± 140.8 vs patient 38063 ± 179.6), (C) rosette circularity (control 0.8789 ± 0.0011 vs patient 0.8749 ± 0.0007), (D) number of lumens per rosette (control 0.9207 ± 0.0403 vs patient 0.7033 ± 0.0209), (E) lumen area (control 5347 ± 211.6 vs patient 6793 ± 105.6), (F) lumen circularity (control 0.7678 ± 0.0053 vs patient 0.7211 ± 0.005), and (G) distance of lumens from the rosette centre (control 23.62 ± 1.15 vs patient 14.6 ± 0.4618). Each data point represents a single rosette. Data were pooled from three independent differentiations and two different patient iPSC clones, with n = 328 control rosettes and n = 718 patient rosettes. Statistical comparisons were performed using a two-tailed Mann–Whitney U test. ****p < 0.0001. Data are shown as mean ± SEM.

Together, these findings demonstrate that *SYNGAP1* haploinsufficiency selectively disrupts neuroepithelial apical organisation at an early developmental stage, prior to the establishment of mature synaptic networks, consistent with an emerging role for *SYNGAP1* in cytoskeletal regulation during neural progenitor development.

### *SYNGAP1* DEE neurons show broad transcriptional dysregulation of neurodevelopmental, cell adhesion and ion channel pathways

To investigate intrinsic transcriptional changes associated with *SYNGAP1* haploinsufficiency, patient and control iPSCs were differentiated into excitatory cortical neurons using a modified NGN2 overexpression protocol ^11^ (Fig. 1A). *SYNGAP1* expression was confirmed by RT-qPCR across the differentiation time course in both patient and control neurons, with progressive upregulation observed in both lines (Fig. 3A). Based on this expression trajectory, DIV 21 and DIV 27 were selected as the primary experimental timepoints for electrophysiology and transcriptional profiling, respectively.

**Figure 3.**
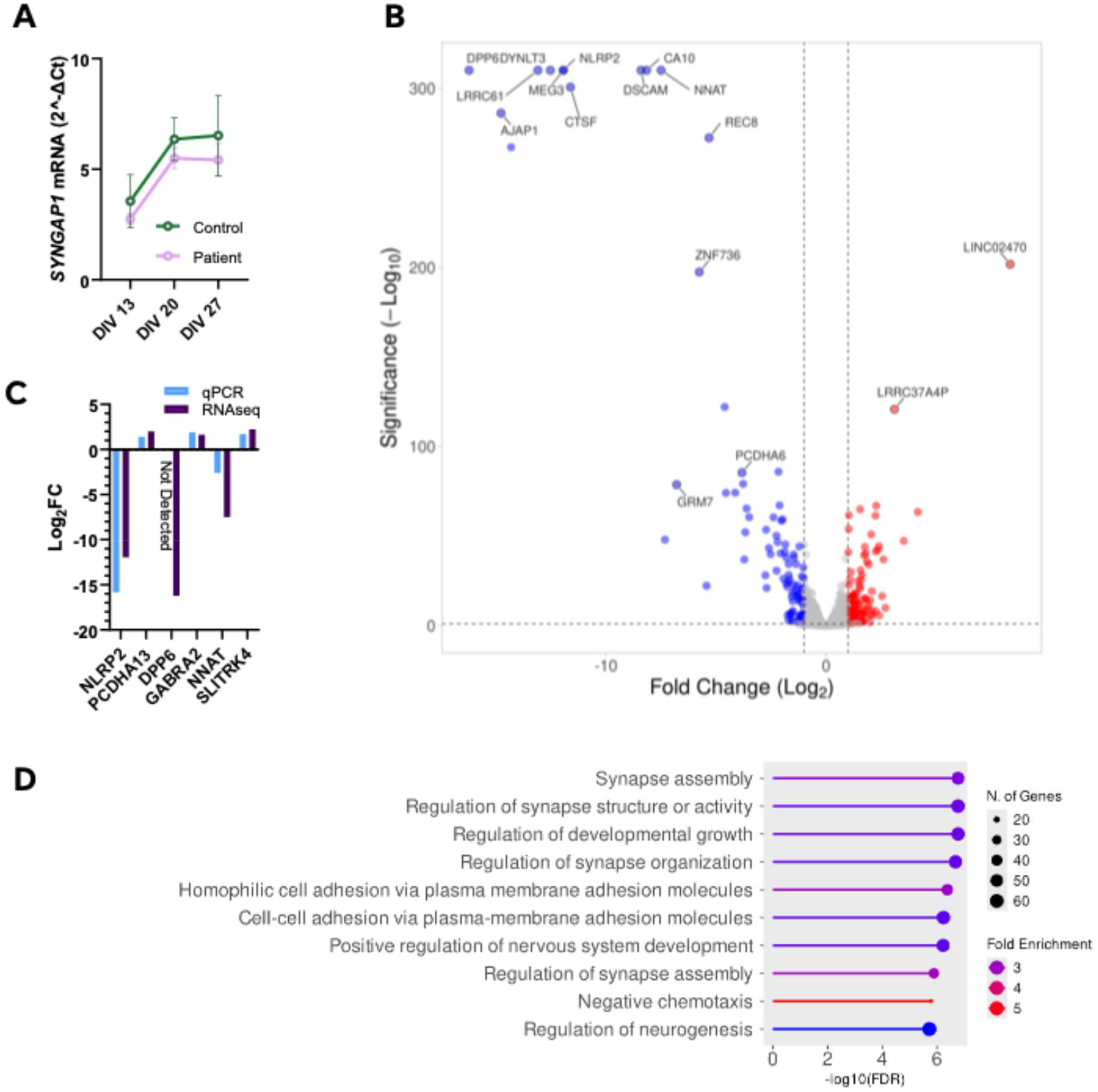
Transcriptomic dysregulation in *SYNGAP1* DEE NGN2-induced neurons at DIV 27. *(A)SYNGAP1* mRNA expression at DIV 13, DIV 20, and DIV 27 in control and *SYNGAP1* DEE cultures. Expression was normalised to GUSB and calculated using the 2^−^ΔCt method. Data are mean ± SEM from n =3 independent differentiations. (B) Volcano plot of differentially expressed genes (DEGs) identified by RNA sequencing of DIV 27 NGN2-induced neurons derived from control and *SYNGAP1* DEE iPSCs. Genes meeting significance criteria (FDR < 0.05) and log_2_ fold change ≥1 are shown in colour (red, upregulated; blue, downregulated); non-significant genes are shown in grey. Selected DEGs are labelled. (C) RT-qPCR validation of selected DEGs. Expression was normalised to GUSB and expressed as log_2_ fold change relative to control neurons. Data are mean ± SEM from n = 3 independent differentiations. (D) Gene Ontology (GO) Biological Process enrichment analysis of DEGs at DIV 27. The top 10 enriched terms are shown ranked by −log_10_(FDR). Dot colour indicates fold enrichment and dot size indicates gene number.

RNA sequencing was performed on three independent differentiation batches of DIV 27 patient and control neurons. Differential gene expression analysis identified 1,929 significantly dysregulated genes (FDR < 0.05) (Fig. 3B). Patient neurons showed downregulation of genes involved in intrinsic neuronal excitability and neurodevelopment, including the voltage-gated potassium channel auxiliary subunit *DPP6* ^12^ and the activity-regulated developmental gene *NNAT* ^13,14^. Among cell adhesion molecules, *DSCAM* and 20 members of protocadherin (PCDH) family were significantly altered (|logFC| > 2, FDR < 0.05). Protocadherins regulate neuronal migration, cell-cell recognition, and circuit assembly, and their disruption has been linked to impaired neural network formation in neurodevelopmental disorders ^15^. The most significantly downregulated gene overall was *NLRP2*, a nucleotide-binding oligomerization domain-containing protein implicated in embryonic brain development and neuropsychiatric disease ^16^.

To validate these findings, RT-qPCR was performed on selected differentially expressed genes in DIV 27 patient and control neurons (Fig. 3C). Directionally consistent changes were confirmed for all validated targets: *PCDHA13, GABRA2*, and *SLITRK4* were upregulated, and *NLRP2* and *NNAT* were downregulated in patient neurons. *DPP6* was undetectable in patient neurons by RT-qPCR, consistent with its pronounced downregulation and large negative log fold change in the sequencing data.

Gene ontology enrichment analysis of differentially expressed genes identified terms predominantly associated with neuronal development and circuit assembly, including regulation of synapse assembly and organisation, homophilic cell-cell adhesion via plasma membrane adhesion molecules, regulation of developmental growth, and nervous system development (Fig. 3D). Enrichment of cell adhesion and synapse assembly terms reflects the widespread dysregulation of protocadherins and *DSCAM*, molecules critical for neuronal identity and wiring specificity during development. Together, these transcriptional findings point to broad dysregulation of neurodevelopmental gene networks in *SYNGAP1* DEE neurons, extending beyond the established synaptic role of *SYNGAP1*.

### *SYNGAP1* neurons exhibit reduced input resistance and larger action potentials

To assess the functional consequences of *SYNGAP1* haploinsufficiency, whole-cell patch-clamp recordings were performed on DIV 20-21 NGN2-induced neurons in the absence of glia, at a stage when these neurons exhibit reliable electrical activity ^17,18,11^.

Analysis of intrinsic membrane properties (Fig. 4A–F) revealed that patient-derived and control neurons had comparable membrane capacitance (Fig. 4A; P = 0.081) and resting membrane potential (Fig. 4B; P = 0.361), indicating similar passive membrane properties. In contrast, *SYNGAP1* DEE neurons displayed significantly reduced input resistance (597.2 ± 70.79 MΩ vs 829.8 ± 66.55 MΩ; Fig. 4C; P = 0.006), consistent with increased membrane conductance. Representative action potential firing traces are shown in Fig. 4D. Patient neurons exhibited a mild rightward shift in the frequency–current relationship at higher stimulus intensities (Fig. 4E), whereas rheobase was comparable between genotypes (Fig. 4F; P = 0.109).

**Figure 4.**
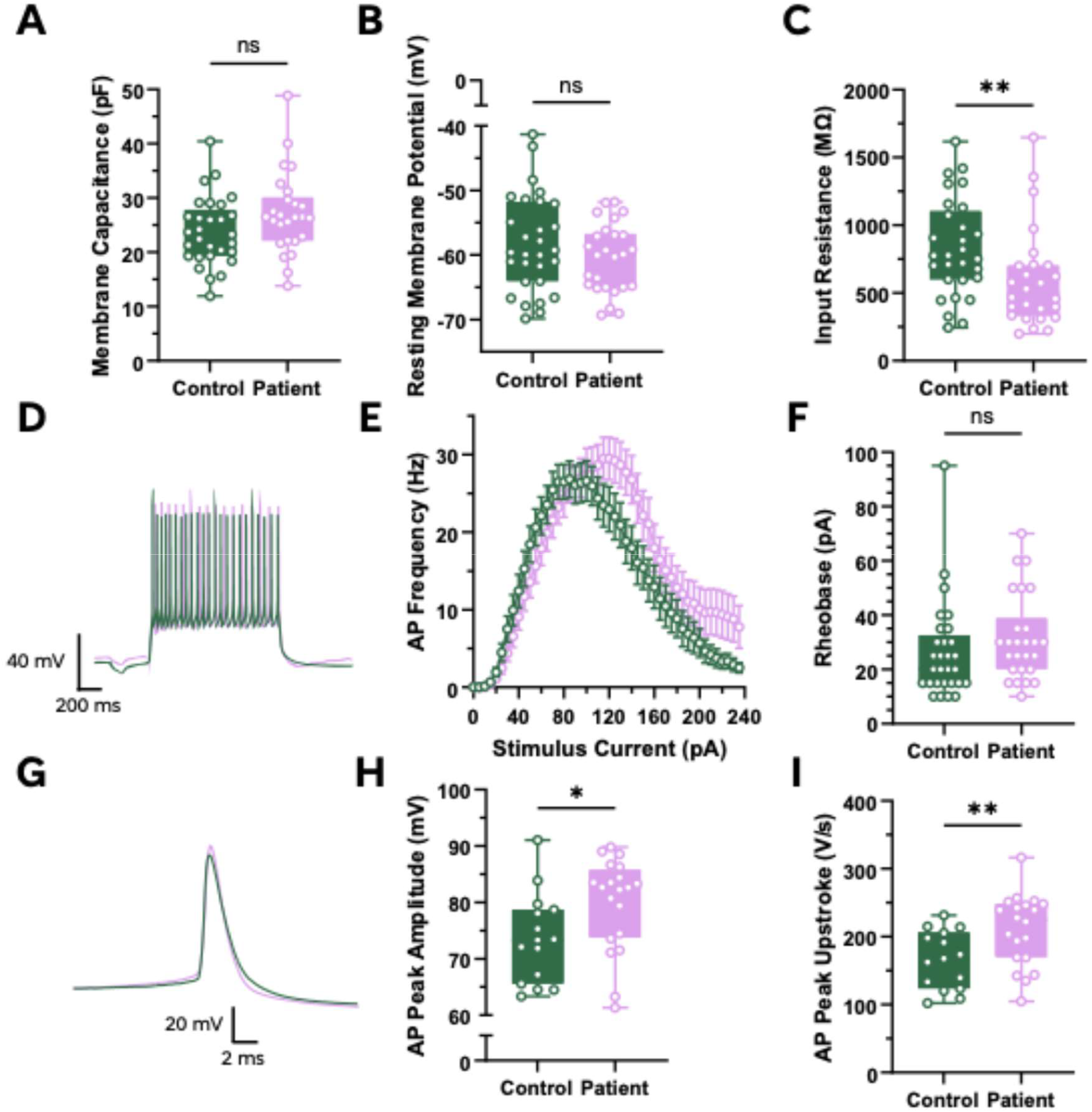
Electrophysiological properties of *SYNGAP1* DEE NGN2 neurons. (A-F) Passive membrane properties and neuronal excitability (n = 29 vs 26 cells, 3 independent differentiations). (A) Membrane capacitance (Ctrl 23.81 ± 1.161 pF vs Patient 27.14 ± 1.457 pF, p = 0.081). (B) Resting membrane potential (Ctrl -58.29 ± 1.387 mV vs Patient -60.48 ± 1.043 mV, p = 0.3611). (C) Input resistance (Ctrl 829.8 ± 66.55 MΩ vs Patient 597.2 ± 70.79 MΩ, p = 0.0056). (D) Representative traces of action potential firing at 60 pA of control (dark green) and patient (pink) neurons. (E) Frequency-Current relationship. (F) Rheobase (Ctrl 26.38 ± 3.291 pA vs Patient 31.54 ± 3.11 pA, p = 0.1091). (G-I) Action potential properties (n = 15 vs 20 cells, 3 independent differentiations). (G) Representative traces of the second action potential at rheobase+1 of control (dark green) and patient (pink) neurons. (H) AP peak amplitude (Ctrl 73.51 ± 2.043 mV vs Patient 79.91 ± 1.822 mV, p = 0.0302). (I) AP peak upstroke (Ctrl 165.8 ± 10.98 V/s vs Patient 210.2 ± 11.74 V/s, p = 0.0085). Data are presented as mean ± S.E.M.

Analysis of action potential waveform properties (Fig. 4G–I) demonstrated significantly larger peak amplitudes (79.91 ± 1.82 mV vs 73.51 ± 2.04 mV; Fig. 4H; P = 0.030) and faster peak upstroke velocity (210.2 ± 11.74 V/s vs 165.8 ± 10.98 V/s; Fig. 4I; P = 0.009) in *SYNGAP1* DEE neurons. Representative action potential traces recorded at rheobase +1 are shown in Fig. 4G.

### *SYNGAP1* DEE neurons have larger current densities

Voltage-clamp recordings were performed to examine inward and outward currents, with the membrane potential held at −70 mV and stepped incrementally (Δ10 mV) to +60 mV (n = 26 vs 24 cells, 3 independent differentiations). Representative inward current traces at −30 mV are shown in Fig. 5A. *SYNGAP1* neurons displayed significantly larger peak inward current density compared to controls (9219 ± 594.5 vs 7256 ± 460.3 pA/pF; Fig. 5B; p = 0.033), with no difference in rise or decay times. Representative outward current traces at +60 mV are shown in Fig. 5C. Steady-state outward current density was also significantly elevated in patient neurons at positive membrane potentials (3427 ± 207.9 vs 2531 ± 88.75 pA/pF; Fig. 5D; p = 0.001).

**Figure 5.**
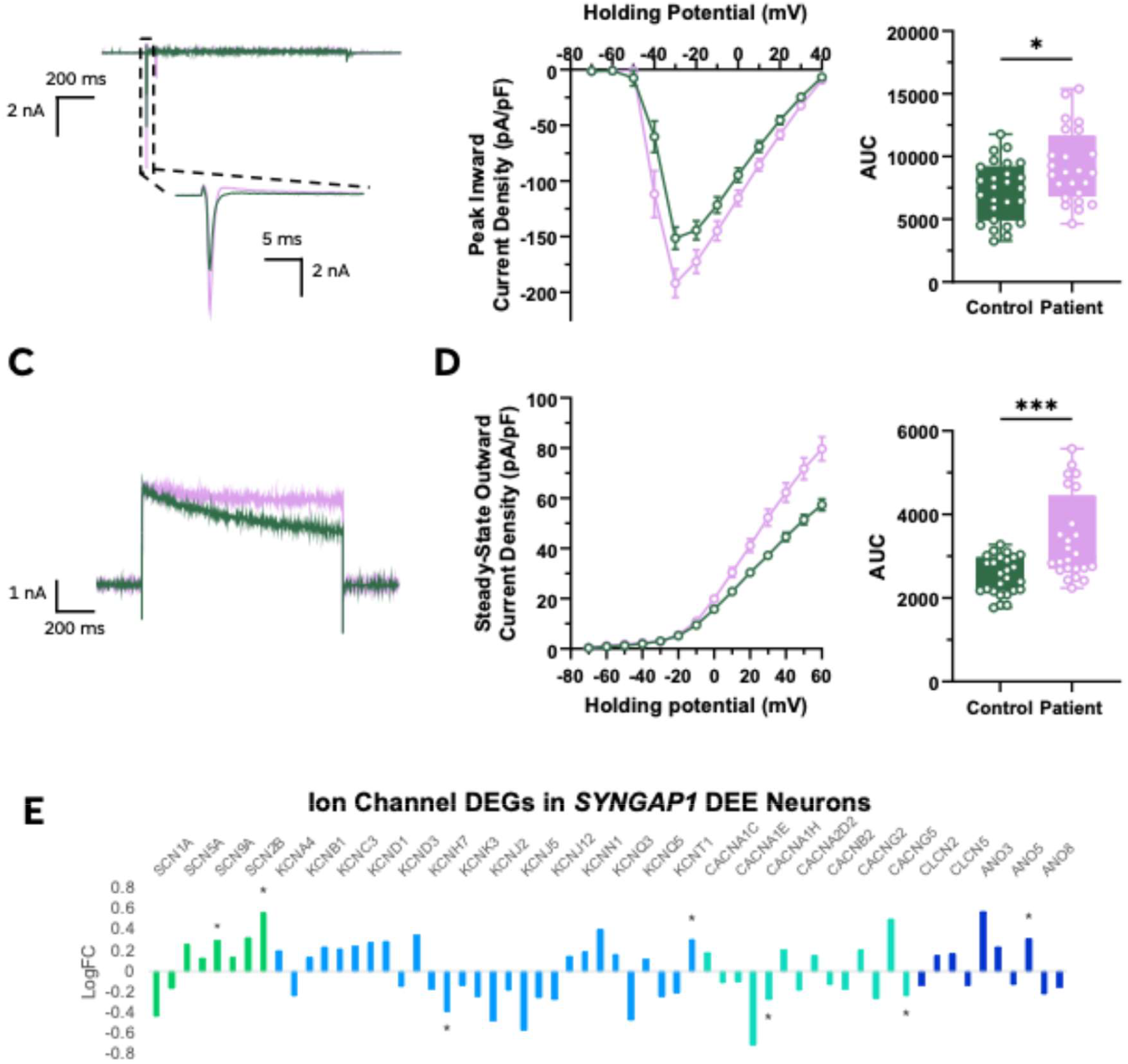
*SYNGAP1* DEE neurons display increased inward and outward current densities. (A) Representative inward currents at -30 mV holding potential including 25 ms window of the peak. (B) Current-Voltage relationship of peak inward current density and area under the curve (Ctrl 7256 ± 460.3 vs Patient 9219 ± 594.5, p = 0.0326). (C) Representative outward currents at +60 mV holding potential. (D) Current-Voltage relationship of steady-state outward current density and area under the curve (Ctrl 2531 ± 88.75 vs Patient 3427 ± 207.9, p = 0.0006). Data are presented as mean ± S.E.M. (n = 26 vs 24 cells, 3 independent differentiations).

## Discussion

This study demonstrates that *SYNGAP1* haploinsufficiency disrupts human neural development across multiple stages, from early neuroepithelial organisation through to the intrinsic functional properties of cortical neurons. Using two complementary patient-derived models, micropatterned neural rosettes and NGN2-induced excitatory neurons, we identify convergent phenotypes that precede the establishment of mature synaptic networks, implicating *SYNGAP1* in neurodevelopmental processes beyond its established role at the post-synaptic density. Collectively, our findings support a model in which early disruption of cytoskeletal organisation, neurodevelopmental gene networks, and intrinsic neuronal maturation contributes to the cortical dysfunction underlying *SYNGAP1* DEE.

The patient line examined here carries c.435_447dup variant resulting in a leucine-to-valine substitution at the duplication junction and a frameshift causing premature protein truncation (p.Leu150Valfs*6). The variant lies within the PH domain, upstream of the RASGAP domain. Given its truncating nature, haploinsufficiency likely results from nonsense-mediated decay of the mutant transcript, leading to loss of the RASGAP domain and other downstream functional regions critical for *SYNGAP1* signalling.

Neural rosettes derived from *SYNGAP1* DEE patient iPSCs displayed consistently enlarged and dysmorphic lumens compared to controls, with reduced lumen circularity and altered lumen positioning relative to the rosette centre. This micropatterned single rosette model recapitulates early neuroepithelial organisation, providing a window into *SYNGAP1* function at a developmental stage preceding neuronal differentiation and synaptogenesis ^9^. Previous study using this platform identified neuroepithelial phenotypes in *SYNGAP1* variants localised to the RASGAP domain ^8^; the present findings extend these observations to a PH domain variant, suggesting that disruption of neuroepithelial architecture is not domain-specific but rather a broader consequence of *SYNGAP1* haploinsufficiency. The observed lumen phenotype points to disrupted apical cytoskeletal organisation in radial glial progenitors, consistent with the known role of *SYNGAP1* in regulating cytoskeletal dynamics through RAS-MAPK signalling ^19,20^. Notably, overall rosette area and circularity were comparable between genotypes, indicating that the structural disruption is specific to apical organisation rather than reflecting a general difference in rosette morphology. The consistency of this phenotype across three independent differentiations and two patient iPSC clones further supports its biological relevance.

To examine whether the early developmental disruption observed in neural rosettes extends to later stages of neuronal differentiation, further analysis was performed on NGN2-induced patient and control neurons. Transcriptomic profiling of NGN2 neurons at DIV 27 revealed widespread dysregulation of genes involved in neurodevelopmental processes, cell adhesion, and ion channel regulation. The most striking finding was the coordinated downregulation of 20 protocadherin family members, predominantly from the alpha cluster, alongside *DSCAM*, molecules that regulate neuronal identity, dendrite morphogenesis, and wiring specificity during cortical development ^15,21^. These findings align with the cytoskeletal disruption observed in rosettes, as alpha-protocadherins regulate cytoskeletal dynamics through the WAVE complex ^21^, suggesting a convergent deficit in cytoskeletal and adhesion machinery across two independent models and developmental stages.

Transcriptomic analysis also identified downregulation of ion channel regulatory genes, including *NNAT* and *DPP6*, which are essential for establishing intrinsic neuronal excitability and maintaining excitatory-inhibitory balance. Among the most significantly downregulated genes overall was *NLRP2*, previously implicated in embryonic brain development and neuropsychiatric disease ^16,22^, though its specific function in human neurons remains poorly understood.

Whole-cell recordings from DIV 20-21 patient neurons revealed a specific electrophysiological signature consistent with accelerated intrinsic maturation rather than generalised hyperexcitability. Input resistance was significantly reduced and action potentials displayed larger peak amplitudes and faster upstroke velocities in patient neurons, while membrane capacitance, resting membrane potential, and rheobase were unchanged. Voltage-clamp recordings demonstrated larger inward and outward current densities, indicative of enhanced sodium and potassium channel function. Critically, the transcriptomic and electrophysiological data are complementary: dysregulation of ion channel regulatory genes may provide a molecular basis for the intrinsic membrane changes, together pointing to accelerated functional maturation as a cell-autonomous consequence of *SYNGAP1* haploinsufficiency.

A plausible mechanistic explanation linking these phenotypes is disruption of RAS-MAPK/ERK signalling, a pathway critically regulated by *SYNGAP1* during cortical development. *SYNGAP1* acts as a RAS GTPase-activating protein, and haploinsufficiency is expected to result in elevated RAS-MAPK/ERK activity through loss of negative regulation ^5,23^. Dysregulated MAPK/ERK signalling during neurodevelopment has well-established consequences for cytoskeletal organisation, neuronal migration, and the transcriptional programmes governing neuronal maturation, processes reflected across all three datasets reported here. The disrupted apical cytoskeletal organisation observed in neural rosettes is consistent with aberrant RAS-MAPK signalling in radial glial progenitors, while the widespread dysregulation of protocadherins, *NNAT, DPP6*, and *NLRP2* in induced neurons suggests broad transcriptional consequences of chronically elevated ERK activity during differentiation. The accelerated intrinsic maturation observed electrophysiologically may similarly reflect MAPK/ERK-driven shifts in ion channel expression and membrane properties during neuronal development. These early cell-autonomous changes likely precede and contribute to later synaptic and network dysfunction, consistent with previous studies in rodent models demonstrating disrupted cortical circuit development and altered GABAergic connectivity in *SYNGAP1* haploinsufficiency ^1,24^. Our findings extend these observations by identifying convergent developmental abnormalities and accelerated intrinsic maturation in human patient-derived models, highlighting the value of human cellular systems for capturing disease-relevant phenotypes.

Several limitations of the present study warrant consideration. First, findings are based on a single patient iPSC line carrying one variant, which limits generalisation across the broader spectrum of *SYNGAP1* pathogenic variants. While two independent patient clones were used for rosette experiments and phenotypes were consistent across three independent differentiations, the absence of an isogenic control line means that genetic background differences between the patient and population control cannot be fully excluded as a contributing factor. Second, NGN2-induced neurons represent a relatively homogeneous population of excitatory cortical neurons and do not recapitulate the cellular diversity of the developing cortex, including inhibitory interneurons and glial populations whose interactions are known to be disrupted in *SYNGAP1* DEE ^1,24^. Third, the transcriptomic and electrophysiological analyses were performed at fixed timepoints, and it remains to be determined whether the phenotypes observed here persist, resolve, or evolve over longer differentiation time courses. These limitations notwithstanding, the convergence of phenotypes across two independent models and multiple levels of analysis strengthens confidence in the biological relevance of the findings reported here.

The present study demonstrates that *SYNGAP1* haploinsufficiency drives disruption of human neural development across multiple stages and biological levels, from neuroepithelial architecture through transcriptional dysregulation to accelerated intrinsic neuronal maturation. These findings support the view that *SYNGAP1* DEE is a disorder of neurodevelopment rather than solely a synaptic pathology, with disease-relevant phenotypes emerging well before mature synaptic networks are established. The identification of convergent cytoskeletal, transcriptional, and functional phenotypes in human patient-derived models provides a mechanistic framework for understanding how early developmental disruption contributes to the cortical dysfunction and intractable epilepsy characteristic of *SYNGAP1* DEE. As gene-targeted therapies for *SYNGAP1* DEE move toward the clinic, these findings highlight the importance of considering neurodevelopmental as well as synaptic mechanisms and therapeutic windows.

## Materials and Methods

### Induced pluripotent stem cells

Two iPSC lines, one patient-derived and one healthy control, were used in this study. The healthy control (Cat #WC026i-5807-3, WiCell, WI, USA) was derived from a healthy female neonate and reprogrammed using episomal based reprogramming factors ^25^. The SYNGAP1 DEE iPSC line was generated from a young female patient skin biopsy. Patient fibroblasts were reprogrammed using episomal based reprogramming factors (cMYC, SOX2, OCT4, KLF4) at StemCore (StemCore, The University of Queensland, QLD, Australia). Detailed methods are described by Ovchinnikov *et al*.^26^.

#### Genotyping

To confirm the presence of the heterozygous *SYNGAP1* variant, specific primers were designed using Primer3Plus (forward primer: CTTTCATCCAGGGGCTCTCTAC; reverse primer: AATCTCATCCCACATGCTCCAG; product length, 241 bp). Final passage genomic DNA from patient and control iPSCs was extracted from a pellet of ∼300000 cells using Qiagen DNeasy Kit (Cat # 69504, Qiagen, Hilden, Germany) according to manufacturer’s instructions. PCR was performed with Taq DNA polymerase (Cat #M02733, New England Biolabs, MS, USA) and amplified the target DNA. The PCR products were cleaned up using PureLink− PCR Purification Kit (Cat #K310001, Thermo Fisher Scientific) and analysed with gel electrophoresis. A total of 100 ng PCR product from control and patient iPSCs was sent to the Australian Genome Research Facility (AGRF, Melbourne, Australia) to undergo Sanger sequencing.

#### Maintenance of iPSCs

All iPSC cultures were maintained in a humidified incubator at 37 °C, 5% CO_2_, and 5% O_2_. Cell lines were routinely tested for mycoplasma using the MycoAlert Detection Kit (Cat #LT07-701, Lonza, Basel, Switzerland) and confirmed to be mycoplasma-free. For routine culture, 6-well plates (Cat #140675, Thermo Fisher Scientific) were coated with vitronectin (Cat #A14700, Thermo Fisher Scientific) diluted in DPBS −/− (Cat #14190144, Thermo Fisher Scientific) for at least 1 h at 37 °C or overnight at 4 °C.

Cryopreserved iPSCs were thawed, centrifuged (300 × g, 3 min), resuspended in Essential 8 (E8) medium (Cat #A1517001, Thermo Fisher Scientific), and plated onto vitronectin-coated wells. Medium was changed daily, and cells were maintained in E8 until 70–80% confluence. Cells were passaged every 3–5 days using 0.5 mM EDTA (Cat #15575020, Thermo Fisher Scientific) at a 1:3– 1:6 split ratio.

For cryopreservation, cells were harvested with EDTA, pelleted (100 × g, 1 min), and resuspended in E8 medium containing 10% dimethyl sulfoxide (DMSO) (Cat #D2650, Sigma-Aldrich, MA, USA). Cryovials were cooled overnight at −80 °C using a Mr. Frosty™ freezing container (Cat #5100-0001, Thermo Fisher Scientific) before long-term storage in liquid nitrogen.

### Neural differentiation

#### Lentiviral Preparation

Lentiviral vectors encoding NGN2 and rtTA were generated in HEK293T cells (Cat #CRL3216, ATCC, VA, USA). HEK293T cells were co-transfected using Lipofectamine™ 3000 Transfection Reagent (Cat #L3000001, Thermo Fisher Scientific) with the lentiviral packaging plasmids pMDL (Addgene #12251), pRSV (Addgene #12253), and pVSV-G (Addgene #8454), together with either mNGN2 (Addgene #52047) or rtTA (Addgene #20342). Plasmid details are provided in Supplementary Table S1.

Following 5–7 h of incubation at 37 °C, transfection medium was replaced with fresh HEK medium. Viral supernatants were collected on days 3, 4 and 5, centrifuged (200 × g, 2 min), filtered through a 0.45 µm syringe filter (Millex-HV, Cat #SLHV033RB, Merck, NJ, USA), and stored at 4 °C until concentration. On day 5, pooled viral supernatants were ultracentrifuged at 25,000 rpm (115,889 × g) for 2 h at 4 °C. Viral pellets were resuspended in 375 µL PBS+/+ (Cat #14040117, Thermo Fisher Scientific), aliquoted, and stored at −80 °C. Each viral preparation was titrated on iPSCs by doxycycline induction and puromycin selection to determine the optimal viral volume for neuronal induction.

#### Generation of NGN2 Neurons

Human induced neurons were generated as previously described (Zhang et al. 2013) with minor modifications. iPSCs were plated at 50,000 cells/well on DIV -2 in E8 supplemented with 10 µM Y27. The following day, DIV -1, fresh E8 medium supplemented with 10 μM Y27 and the two lentivirus particles, *NGN2* and rtTA, was transfected into the iPSCs. Cells were cultured in a low oxygen incubator overnight for 16-20 hours. *NGN2* was induced by the addition of doxycyxline to culture medium. From DIV 1 to 3, cells underwent Puromycin selection. On DIV 4, cells were lifted and dissociated to be replated onto 24 well plates coated with poly-D-lysine (100 μg/ml in sterile water) (Cat #P7280, Sigma Aldrich) and Laminin 2020 (15 µg/ml).

For RNA experiments cells were plated at 100,000 cells per well in 500 µl media, while for patch clamping and immunocytochemistry, cells were plated at 50,000 per well in 500 µl media on plates with glass coverslips (Cat #6302118, Thermo Fisher Scientific).

#### Neuron and Astrocyte Co-cultures

Human fetal astrocytes were obtained from ScienCell Research Laboratories (1801) and cultured according to manufacturer’s instructions.

### RNA isolation and qRT-PCR

To quantify *SYNGAP1* gene expression in patient and control neurons, real-time quantitative PCR was performed at three timepoints of the NGN2 neuron protocol. A primer targeting exon 8-9 of the *SYNGAP1* gene was used to detect the expression levels.

Three biological replicates of NGN2 control and patient neurons were collected, and total RNA was isolated using the Purelink RNA Mini Kit. 4-6 wells of neurons were harvested at 3 timepoints, DIV 13, DIV 20, and DIV 27. The PureLink Mini Kit was used according to manufacturer’s instructions. Samples were eluted using 30 µl RNase free water. Sample concentration and purification were quantified using the Nanodrop 2000 Spectrophotometer (Cat #ND-2000, Thermo Fisher Scientific. The purified RNA was stored at -80 °C.

Real time quantitative PCR was conducted on the QuantStudio 7 in combination with the Taqman RNA-to-Ct 1-step Kit (Cat #4392653, Thermo Fisher Scientific) according to the manufacturer’s directions. Real time qPCR experiments were conducted in duplex reactions, in triplicate, to obtain the relative levels of each transcript normalized to the housekeeping gene *GUSB* within a single well. For each sample, the cycle threshold (Ct) values from three technical replicates were averaged. Each reaction contained 32 ng of RNA and the total reaction volume was 10 µl.

### Electrophysiology

Whole-cell patch clamping was performed on DIV 20-21 neurons from three differentiations, with patient and control lines recorded in parallel. Glass coverslips were transferred to a 0.3 ml recording chamber superfused with BrainPhys™ Neuronal Medium (Cat #05790, Stem Cell Technologies, Australia). Cells were patched with thick-walled glass patch pipettes (Warner Instruments, USA) at 30 ± 1°C with a flow rate of 1 ml/min, using a pipette resistance of 3-5 MΩ. Patch pipettes were filled with potassium gluconate-based intracellular solution (in mM): 130 K-gluconate, 10 D-glucose, 6 KCl, 5 EGTA, 5 HEPES, 4 NaCl, *2* MgATP, *and 0*.*3* GTP-tris salt (pH adjusted to 7.3 with KOH, 290 mOsm). Cells were excluded from recordings if their resting membrane potential was >-40 mV, membrane capacitance <12 pF or if the access resistance >30 MΩ.

Spontaneous activity and the resting membrane potential were measured using I=0 (gap-free) immediately after establishing the whole-cell patch clamp configuration. The cell’s resting state was recorded for a minimum of 45 seconds followed by an episodic current clamp protocol that involved holding the cells at -70 mV between current steps and injecting them with increasing amounts of current from -10 to 235 pA at 5 pA intervals, spanning 1000 ms each. Lastly, inward and outward currents were measured using an episodic voltage clamp protocol. The cells were held at -70 mV between voltage steps and membrane potential was gradually increased from -70 to 40 mV at 10 mV intervals spanning 500 ms each. Action potential threshold was taken as the membrane potential at which dv/dt = 20 V/s.

Recordings were digitised at 10 kHz and filtered off-line at 5 kHz. Neurons meeting the following criteria were used: series resistance <30 MΩ, resting potential < -40 mV, and holding current > - 100 pA. Input resistance was calculated as the slope of the voltage-current curve determined using three 1000 ms current steps (Δ5 pA) from -10 pA to 0 pA. For the analysis of action potential (AP) morphology, the second AP from the first current step above rheobase was used. AP threshold was calculated where the first derivative of the up phase of the trace equalled 20 V/s. The rise and decay times were calculated from 10-90% of the peak value and the half-height width was measured at 50% of the peak amplitude.

Whole-cell patch clamp data was acquired using Multiclamp 700B controlled by Clampex 10.7 / Digidata 1550B acquisition systems (Molecular Devices, USA). Data analysis was done with Clampfit 10.7 (Molecular Devices, USA) and GraphPad Prism 10 (GraphPad, USA).

### RNA Sequencing

NGN2 neuron samples were processed for RNA sequencing using the PureLink RNA Mini Kit™ as detailed above. Neurons from two 24-well plates were collected from three biological replicates of control and patient cell lines. RNA samples underwent DNase treatment using DNA-*free* DNA removal Kit (Cat #AM1906, Thermo Fisher Scientific) and quantified using the NanoDrop 2000. RNA sequencing was performed by the Australian Genome Research Facility (AGRF; Melbourne, Australia) using the Illumina NovaSeq platform with stranded poly(A)-selected mRNA libraries, generating approximately 150 million paired-end reads (150 bp) per sample.

mRNA-seq read quality was assessed using the FastQC software version 0.11.9 (www.bioinformatics.babraham.ac.uk/projects/fastqc/). Alignment and quantification of mRNA-seq data was performed using Rsubread aligner (version 2.4.3). Paired-end 150bp Illumina reads were aligned to the human genome (Ensembl GRCh38 assembly) at the gene-level with an average of 96.47% (± 0.19 SD) of reads successfully mapped. Genes with greater than five counts-per-million mapped reads in at least two samples were retained for further analysis, genes below this threshold were filtered.

Differential gene expression analysis was performed in the R statistical programming environment (version 4.0.5) using EdgeR ^27^ (version 3.32.1), which applied adjustment for library size and normalisation with trimmed mean of M values (TMM) where dispersion parameters for each gene are estimated with the Cox-Reid common dispersion method and employed in a negative binomial generalized linear model for each gene. Accounting for gene dispersion ensures that expression differences that are consistent between replicates are more highly weighted than those that are not to ensure differential expression is not driven by outliers. P-values were adjusted for multiple testing using the Benjamini-Hochberg correction with an FDR q-value <0.05 cut off.

Gene ontology enrichment analysis performed on the differential expressed genes was conducted using Gorilla (http://cbl-gorilla.cs.technion.ac.il/)^28,29^. The GO terms were summarised and visualised by uploading to REVIGO (http://revigo.irb.hr/.) where terms are grouped by semantic similarities ^30^.

### Neural Rosettes

#### Differentiation of Neural Rosettes

Human iPSCs were differentiated into neural rosettes using RosetteArray™ 96-well plates. Plates were rinsed once with PBS−/− and coated with growth factor-reduced Matrigel diluted in cold Essential 8™ medium (Gibco, A1517001). One hundred microlitres of Matrigel solution was added per well and incubated at 37 °C for 18–20 h.

iPSCs were dissociated with Accutase, centrifuged at 300 × *g* for 3 min, and resuspended in Essential 8™ medium containing 10 µM Y-27632 ROCK inhibitor. Cells were seeded at 64,000 cells per well. After overnight incubation, neural induction was initiated by replacing 75% of the medium with Essential 6™ medium (Gibco, A1516401). From DIV 1–6, 50% medium changes were performed daily. On DIV 7, cultures were fixed with 4% paraformaldehyde for 20 min at 4 °C, washed twice with PBS−/−, and stored in PBS at 4 °C.

#### Immunocytochemistry

Fixed rosettes were blocked and permeabilised in PBS containing 10% heat-inactivated fetal bovine serum and 0.2% Triton X-100 for ≥1 h. Primary antibodies were applied overnight at 4 °C: rabbit anti-PAX6 (1:250; BioLegend, 901301) and mouse anti-N-cadherin (1:500; BD Biosciences, 610920). After PBS washes, secondary antibodies were applied for 1 h at room temperature: goat anti-rabbit Alexa Fluor 594 (1:1000; Invitrogen, A32740) and goat anti-mouse Alexa Fluor 488 (1:1000; Invitrogen, A11001). Nuclei were stained with DAPI (1:1000) for 5 min and wells were sealed with ProLong™ Glass Antifade Mountant (Invitrogen) before imaging.

#### Imaging of Neural Rosettes

Neural rosettes were imaged using a Nikon Ti2-E inverted microscope stand equipped with a CrestOptics X-light V3 spinning disk confocal with 50 µm pinholes. Imaging for analysis was performed using a Plan Apo 20×/0.8NA (Nikon) objective. Fluorophores were excited using 405 nm, 488 nm, and 555 nm lasers, (89 North LDI-5) and 440/40, 525/30 and 600/30 filter sets respectively. Each laser was set to 80% power with 300ms exposure times. A Photometrics Kinetix sCMOS camera was used for image acquisition at a resolution of 3200 × 3200 pixels and 12-bit depth.

Z-stacks were collected using a piezo stage (Mad City Labs), spanning a total range of 10 µm centred on the neural rosette mid-plane (±5 µm) with a step size of 0.5 µm, yielding 21 optical sections per stack

Images for cell counting and representative high-resolution images were acquired using a Plan Apo Lambda S 40×/1.25NA silicone oil immersion (Nikon). Images were collected at 2048 × 2048-pixel resolution. Z-stacks were obtained over a 5 µm range centered in the middle of the rosette (±2.5 µm), with 0.5 µm steps.

#### Image Analysis

Neural rosette and lumen boundaries were annotated on maximum-intensity projections of the N-cadherin channel using a semi-automated approach implemented in Labelme (v5.4.1; Kentaro Wada, GitHub), which uses the EfficientSAM AI model for assisted segmentation. In cases where lumina appeared faint and EfficientSAM was unable to generate reliable segmentations, regions of interest (ROIs) were manually delineated. Morphological features were then extracted from the resulting segmentations using a custom Python analysis pipeline, including rosette area, lumen area, circularity, the distance between the centres of mass of the lumen and rosette, and the number of lumina per rosette.

### Statistical Analysis

Statistical analyses were performed using GraphPad Prism 9.2. Data are presented as mean ± SEM. Statistical tests used for each analysis are indicated in the corresponding figure legends. P < 0.05 was considered statistically significant.

## Declarations

### Author contributions

M.W. and L.T. conceived and performed cell differentiation, experimental work, data collection and analysis, and contributed equally to manuscript writing. M.W. led iPSC culture, neural differentiation and neural rosette experiments. L.T. performed electrophysiological experiments and contributed to data analysis. S.B. performed RNA sequencing bioinformatic analyses. N.E.R. and H.P. contributed to neural rosette image analysis. D.O. generated the *SYNGAP1* patient iPSC line (NGF021). C.M. assisted with experimental work and manuscript preparation. I.E.S. recruited patients, obtained fibroblasts and provided clinical data. S.P. contributed to study conception, experimental design and structural support. S.M. conceived and supervised the study, secured funding, and wrote and revised the manuscript. All authors reviewed and approved the final manuscript.

### Data availability statement

RNA sequencing data generated in this study will be deposited in the Gene Expression Omnibus (GEO) and the accession number provided prior to publication. All other data supporting the findings of this study are available from the corresponding author upon reasonable request.

### Code availability

Any custom code used in this study is available from the corresponding author upon reasonable request.

## Acknowledgements

We sincerely thank the patient and her family for participating in this study and for their ongoing commitment to advancing *SYNGAP1* research. We also gratefully acknowledge the support of the SYNGAP Research Fund Australia, administered through the Epilepsy Foundation, for supporting this work.

## Ethics Approval

This study was approved by the Austin Health Ethics committee, ethics number HREC/16/Austin/472.

