## Supplementary for "*SYNGAP1* haploinsufficiency disrupts early neurodevelopment and accelerates intrinsic neuronal maturation in human patient-derived models"

* These authors contributed equally

### Contents

- **Supplementary Figure S1.** Quality control and characterisation of control and SYNGAP1 patient-derived iPSC lines
- **Supplementary Table S1.** Lentiviral plasmids used for NGN2 neuronal differentiation

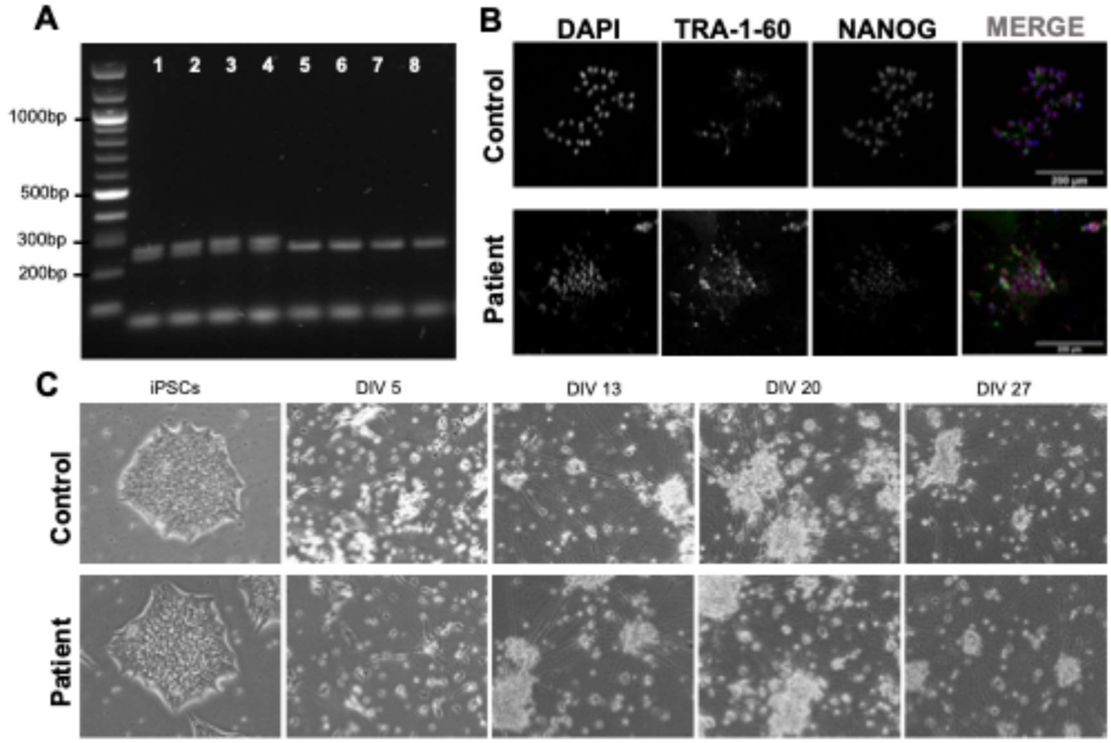


**Supplementary Figure S1. Quality control and characterisation of control and SYNGAP1 patient-derived iPSC lines.** (A) PCR amplification across the SYNGAP1 c.435_447dup variant confirms heterozygous carriage of the duplication in patient-derived iPSCs (lanes 1-4), while the control line produces the expected wild-type amplicon (lanes 5-8). (B) Immunocytochemical staining demonstrates expression of the pluripotency markers TRA-1-60 and NANOG in both control and patient iPSC lines. Nuclei were counterstained with DAPI. (C) Representative brightfield images showing control and patient iPSCs and NGN2-induced neuronal differentiation at the indicated time points (iPSCs, DIV5, DIV13, DIV20 and DIV27), demonstrating comparable neuronal differentiation and culture morphology.

**Supplementary Table S1. Plasmid Details for Lentivirus Production**

| Plasmid | Contents | Addgene # | μg/flask |
|---|---|---|---|
| **Ngn2 [23]** | Mouse neurogenin-2 and puromycin resistance gene under a TetON promoter | 52047 | 12 |
| **pMDL [24]** | Packaging plasmid (Gag and Pol) | 12251 | 6 |
| **pRSV [24]** | Packaging plasmid | 12253 | 3 |
| **pVSV [25]** | Envelope protein | 8454 | 3 |
| **rtTA [26]** | TetON reverse tetracycline transactivator | 20342 | 12 |
